# Evaluation of a scalable approach to generate cell-type specific transcriptomic profiles of mesenchymal lineage cells

**DOI:** 10.1101/2022.12.13.520148

**Authors:** Luke J Dillard, Will T Rosenow, Gina M Calabrese, Larry D Mesner, Basel M Al-Barghouthi, Abdullah Abood, Emily A Farber, Suna Onengut-Gumuscu, Steven M Tommasini, Mark A Horowitz, Clifford J Rosen, Lutian Yao, Ling Qin, Charles R Farber

## Abstract

Genome-wide association studies (GWASs) have revolutionized our understanding of the genetics of complex diseases, such as osteoporosis; however, the challenge has been converting associations to causal genes. Studies have demonstrated the utility of transcriptomics data in linking disease-associated variants to genes; though for osteoporosis, few population transcriptomics datasets have been generated on bone or bone cells, and an even smaller number have profiled individual cell-types. To begin to evaluate approaches to address this challenge, we profiled the transcriptomes of bone marrow-derived stromal cells (BMSCs) cultured under osteogenic conditions, a popular model of osteoblast differentiation and activity, from five Diversity Outbred (DO) mice using single-cell RNA-seq (scRNA-seq). The goal of the study was to determine if BMSCs could serve as a model for the generation of cell-type specific transcriptomic profiles of mesenchymal lineage cells derived from large populations of mice to inform genetic studies. We demonstrate that dissociation of BMSCs from a heavily mineralized matrix had little effect on viability or their transcriptomic signatures. Furthermore, we show that BMSCs cultured under osteogenic conditions are diverse and consist of cells with characteristics of mesenchymal progenitors, marrow adipogenic lineage precursors (MALPs), osteoblasts, osteocyte-like cells, and immune cells. Importantly, all cells were nearly identical from a transcriptomic perspective to cells isolated directly from bone. We also demonstrated the ability to multiplex single cells and subsequently assign cells to their “mouse-of-origin” using demultiplexing approaches based on genotypes inferred from coding SNPs. We employed scRNA-seq analytical tools to confirm the biological identity of profiled cell-types. SCENIC was used to reconstruct gene regulatory networks (GRNs) and we showed that identified cell-types show GRNs expected of osteogenic and pre-adipogenic lineage cells. Further, CELLECT analysis showed that osteoblasts, osteocyte-like cells, and MALPs captured a significant component of BMD heritability. Together, these data suggest that BMSCs cultured under osteogenic conditions coupled with scRNA-seq can be used as a scalable and biologically informative model to generate cell-type specific transcriptomic profiles of mesenchymal lineage cells in large mouse, and potentially human, populations.

## Introduction

Osteoporosis is a disease characterized by low bone mineral density (BMD) and an increased risk of fracture^1^. Osteoporosis-related quantitative traits, such as BMD, are highly heritable^2^ and genome-wide association studies (GWASs) for BMD have identified over 1,100 independent associations^3^. The goal of BMD GWAS is to identify the responsible causal genes^4,5^. However, this is often difficult due to challenges, such as linkage disequilibrium between potentially causal variants^5^ and the observation that most associations implicate non-coding variation^6^. The generation of transcriptomics data and use of systems genetics approaches to interpret GWAS can address these limitations by assisting in prioritizing putatively-causal genes for further investigation^7,8^.

The utility of transcriptomic data to inform BMD GWAS has been demonstrated through studies using approaches such as expression quantitative trait locus (eQTL) mapping and colocalization^9–11^, transcriptome-wide association studies (TWASs)^12,13^, and reconstruction of transcriptomic networks (e.g., gene-regulatory and co-expression networks)^14–16^. These studies have utilized bone, non-bone (e.g., the Gene Tissue Expression (GTEx) project)^17^, and mouse bone transcriptomic data. However, all of the transcriptomic data used to inform BMD GWAS to date has been generated using bulk RNA-seq. These samples are a mixture of data derived from all cells associated with a particular microenvironment and downstream data analysis is often constrained by the inability to definitively attribute transcriptomic signatures to a single cell-type^18^. Further, signals from potentially rare cell-types can be masked by the presence of more abundant cell populations^19^. As a result, there is currently a need to generate population-scale (i.e., hundreds of samples) cell-type specific expression data on cells directly relevant to bone to aid in the identification of causal BMD GWAS genes.

In recent years, single-cell RNA-seq (scRNA-seq) has enabled the efficient generation of high-quality transcriptomes in individual cells^20^. ScRNA-seq can remedy the aforementioned challenges posed with bulk RNA-seq by enabling the generation of single-cell transcriptomic profiles from heterogeneous tissues or primary cell cultures. ScRNA-seq has provided significant insight into the landscape of bone cell-types^21–25^. However, we still lack cost-effective approaches capable of generating scRNA-seq data at scale for key bone cell-types.

Here, we explored the use of bone-marrow derived stromal cells (BMSCs) cultured under osteogenic conditions (BMSC-OBs), a popular *in vitro* model of osteoblast differentiation, to address the above limitations by generating scRNA-seq data on cells of the mesenchymal lineage. We sought to explore technical challenges, cellular heterogeneity, and compare cultured cells to the same cells isolated directly from bone. We show that this approach not only enriches for osteogenic cells, but is a scalable approach capable of generating biologically informative cell-type specific transcriptomic profiles relevant to BMD GWAS. Our results suggest that scRNA-seq of BMSC-OBs has the potential to enable the large-scale generation of cell-type specific transcriptomic data on mesenchymal lineage cells that can be used to inform genetic studies in mice and potentially humans.

## Results

### BMSC cultures grown under osteogenic differentiation conditions are heterogenous

We isolated BMSCs from five Diversity Outbred (DO) mice (N=4 males and N=1 female). The DO is a genetically diverse outbred population derived from eight inbred laboratory strains^26^. We have previously used the DO to perform GWAS for bone strength traits^16^. BMSCs were cultured under osteogenic conditions for 10 days and demonstrated mineralized nodules as previously shown in ^16^. After differentiation, cells were liberated from mineralized cultures and profiled using scRNA-seq. After stringent pre-processing and quality control of the data (**Methods**), 17,457 genes were identified in 7357 cells across all five mice. Unsupervised clustering identified eight distinct cell clusters ranging in size from 46 to 2367 cells (**Figure 1A**).

**Figure 1:**
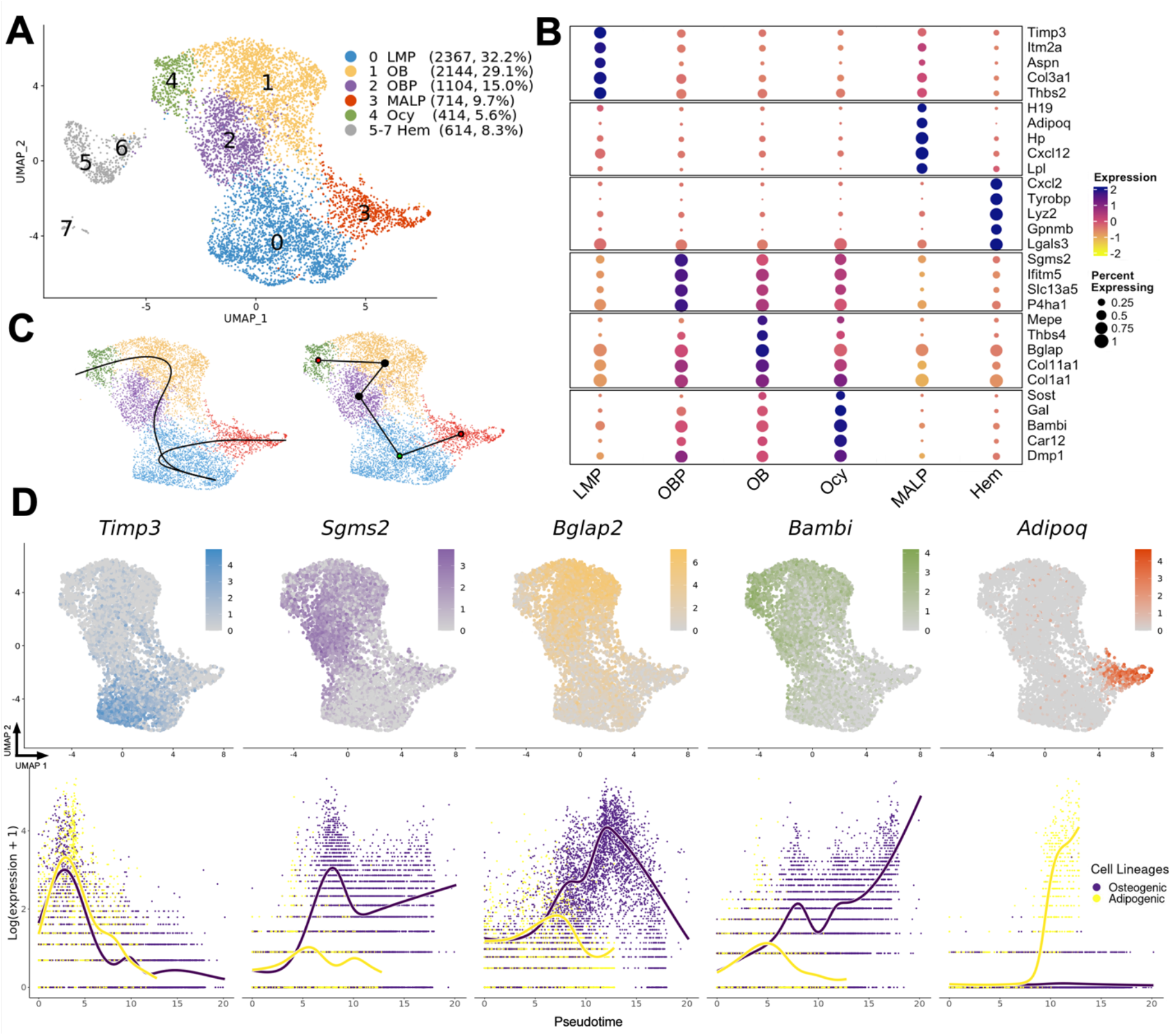
ScRNA-seq of BMSC-OBs identifies multiple cell-types. **A)** UMAP cell clusters of 7357 single BMSC-OBs isolated from five Diversity Outbred (DO) mice. Cell numbers and corresponding percentages are listed in parenthesis to the right of the annotated cluster name. LMP: late mesenchymal progenitor cells; MALP: marrow adipogenic lineage precursors; OBP: osteoblast progenitor cells; OB: osteoblasts; Ocy: osteocytes; Hem: Hematopoietic lineage cells. **B)** Dot plot of some of the most highly expressed genes for all annotated cell clusters. The size of the dots are proportional to the percentage of cells of a given cluster that express a given gene while the color of the dot corresponds to the normalized expression. **C)** Slingshot trajectory inference plots portraying bifurcating branched lineages deriving from LMPs to their respective osteogenic (Ocy) or adipogenic (MALPs) cell fates represented as smooth curves (left plot) or dotted line (right plot). Starting cluster (LMP) is indicated by a green dot and terminal cell fates of the lineages (Ocy, MALPs) are red (right plot). **D)** Feature plots portraying the normalized expression of select genes associated with each cell cluster (top) and each gene plotted as a function of pseudotime overlaid with cell lineages (osteogenic and adipogenic).

We manually annotated the cell-type identity of each cluster using the “FindAllMarkers” function in Seurat^27^ to highlight differentially expressed genes (DEGs) for each cluster relative to all other clusters (**Supplemental Data 1**). As a framework, we used the nomenclature of Zhong et al. (2020) who labeled, isolated, and sequenced single cells from bone marrow using *Col2-Cre Rosa26 <lsl-tdTomato>* reporter mice^23^. In these mice, tdTomato (Td) labels cells spanning the mesenchymal lineage. From Td+ cells, three types of mesenchymal progenitors were identified: early (EMPs), intermediate (IMPs), and late (LMPs). None of the BMSC-OB clusters reported here had signatures of EMPs or IMPs (**Figure 1A**); however, cluster 0 (32.2% of the cells) had high expression of marker genes associated with LMPs, such as *Aspn, Timp3, Thbs2*, and *Itm2a* (**Figure 1A, 1B**). Clusters 1, 2, and 4, (49.7% of the cells) all had signatures of cells in the osteoblast lineage. Mature osteoblasts (Cluster 1) exhibited expression of *Bglap* and *Mepe*, while Cluster 4 had a transcriptomic signature of osteocyte-like cells with high expression of *Bambi* and *Sost* (**Figure 1A, 1B**). Cells in Cluster 2 resembled an osteoblast progenitor (OBP) population differentiating into mature osteoblasts and expressed genes such as *Sgms2, Ifitm5*, and *P4ha1* (**Figure 1A, 1B**). Marrow adipogenic lineage precursors (MALPs), identified as a novel component of bone marrow in Zhong et al. (2020) were represented in Cluster 3 (accounting for 9.7% of the cells) and expressed known MALP markers (*Cxcl12, Adipoq, H19, Hp, Lpl*) (**Figure 1A, 1B**). Cluster 5, 6, and 7 (8.3% of the cells) were cells not associated with the mesenchymal cell lineage and have transcriptomic signatures of immune cells derived from the hematopoietic cell lineage (**Figure 1A, 1B**). Trajectory analysis (cell lineage and pseudotime inference) using Slingshot^28^ on the mesenchymal lineages revealed the expected bifurcating lineage relationship in which LMPs give rise to MALPs and osteoblast progenitors/osteoblasts, independently, with osteocyte-like cells downstream of osteoblasts (**Figure 1C**). The expression of marker genes representative of all cell-types as a function of pseudotime were consistent with cell-type annotations (**Figure 1D**).

### Cell clustering is robust to the effects of cell isolation

The isolation of cells from their heavily mineralized matrix (as outlined in **Methods**) took approximately two hours, raising the possibility that the procedure itself could have an effect on gene expression, transcriptomic signatures, and downstream clustering of cells. To directly assess the effects of the single-cell isolation procedure, we performed a separate experiment in which we generated two identical cultures of BMSC-OBs (10-days post differentiation as in the scRNA-seq experiment) from C57BL/6J mice (N=7) (**Figure 2A**). From one culture (**bulk**), we extracted RNA from the entire well and performed RNA-seq. From the other culture (pooled single cell-bulk, **psc-bulk**), cells were harvested via the single-cell isolation procedure, pooled into one sample, and profiled using RNA-seq. Overall gene expression between the bulk and psc-bulk samples was highly correlated (r=0.98, P<2.2 × 10^−16^) (**Figure 2B**). However, a total of 776 genes were differentially expressed (P_adj_<0.05) with a fold-change less than 0.5 and greater than 2.0 in the psc-bulk vs. bulk samples (**Supplemental Data 2**). A PANTHER^29^ Gene Ontology (GO) enrichment analysis revealed that differentially expressed genes (DEGs) consisted of “acute inflammatory response” (GO:0002675, N =11, P=2.43 × 10^−8^) and “response to stress” (GO:0080134, N = 111, P= 4.96 × 10^−19^) signatures (**Supplemental Data 3)**. To evaluate the impact of the single-cell isolation procedures on cell clustering of the scRNA-seq dataset, we removed the psc-bulk DEGs from the scRNA-seq count matrix. Of the 776 DEGs, 703 (91%) were also captured in the scRNA-seq dataset. Upon removal, a negligible effect was observed on the cell clustering in UMAP space and six distinct cell clusters (five mesenchymal lineage cell clusters) were annotated, similar to the original UMAP (**Figure 2C**). Only 8.1% of cells shifted cell cluster assignment upon removal of DEGs (**Figure 2D**). Most of the cells with shifted assignments were located on the boundaries of cell clusters (**Figure 2D**). These data indicate that gene expression is altered in a predictable manner by the cell isolation procedure, but has little meaningful impact on cell clustering.

**Figure 2:**
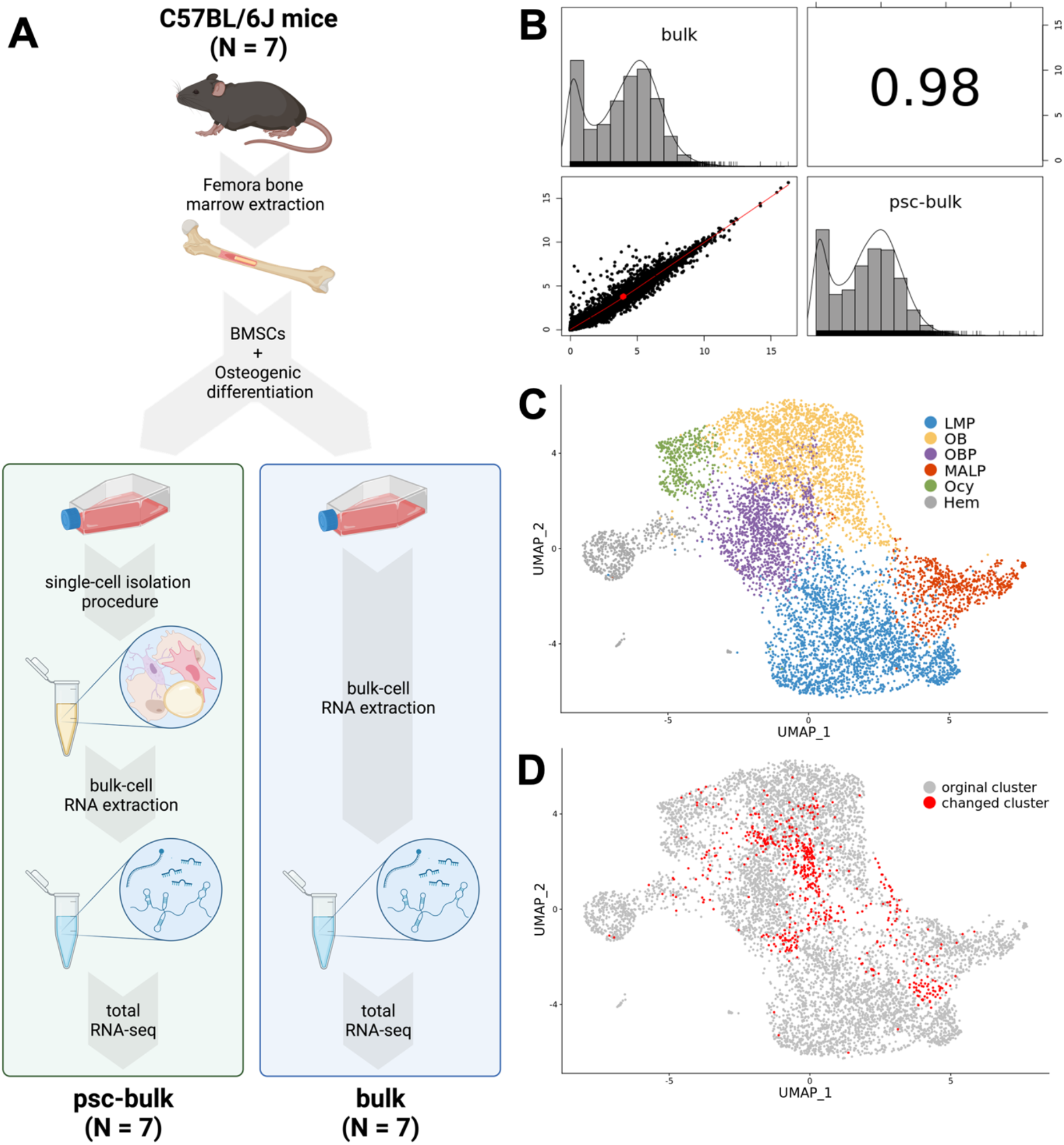
Liberation of single cells from a heavily mineralized matrix *in vitro* has minimal impact on transcriptomic signatures of BMSC-OBs. **A)** Flow chart diagram portraying the design of the bulk vs. psc-bulk experiment in C57BL/6J mice (N=7). Cultured BMSC-OBs were harvested and underwent either immediate RNA extraction (bulk) or the single-cell isolation procedure, pooled, and then subsequent RNA extraction (psc-bulk). Extracted RNA from both conditions was sequenced via traditional RNA-seq. Created with BioRender.com. **B)** Correlation (r=0.98, P<2.2 × 10^−16^) between the counts per million (CPM) values derived from RNA-seq counts for bulk and psc-bulk samples. **C)** scRNA-seq UMAP clusters of BMSC-OBs derived from the five DO mice after removal of differentially expressed genes (identified from the psc-bulk vs. bulk experiment, 703 total genes) from the scRNA-seq count matrix. **D)** Cells highlighted in red represent those that changed from their original cell cluster annotation as a result of removal of DEGs (8.1% of cells).

### Cell-types isolated from BMSC-OBs are similar to their in vivo counterparts

We next wanted to determine if mesenchymal cells generated *in vitro* were similar, in terms of global gene expression, to cell-types isolated directly from bone. Zhong et al. (2020) performed scRNA-seq on Td+ bone marrow cells from mice at 1, 1.5, and 3 months of age^23^. We jointly processed the data from both experiments and integrated the datasets using Canonical Correlation Analysis (CCA)^30^. Overall, the cells from both experiments displayed significant overlap (**Figure 3A**). This was even more apparent when clusters were annotated and cell-types (LMPs, MALPs, OBs, and Ocy-like cells) overlapped in UMAP space between the datasets (**Figures 3B**). A notable difference between cell-types was the absence of EMPs and IMPs in the cultured BMSC-OBs. However, an appreciable enrichment of osteoblast lineage cells, particularly in the OBP population, was observed in the BMSC-OB data compared to the cells isolated directly from bone (**Figure 3B, 3C**). Importantly, the overlap of cells from the two studies suggests few transcriptional differences as a consequence of cell culture and *in vitro* differentiation.

**Figure 3:**
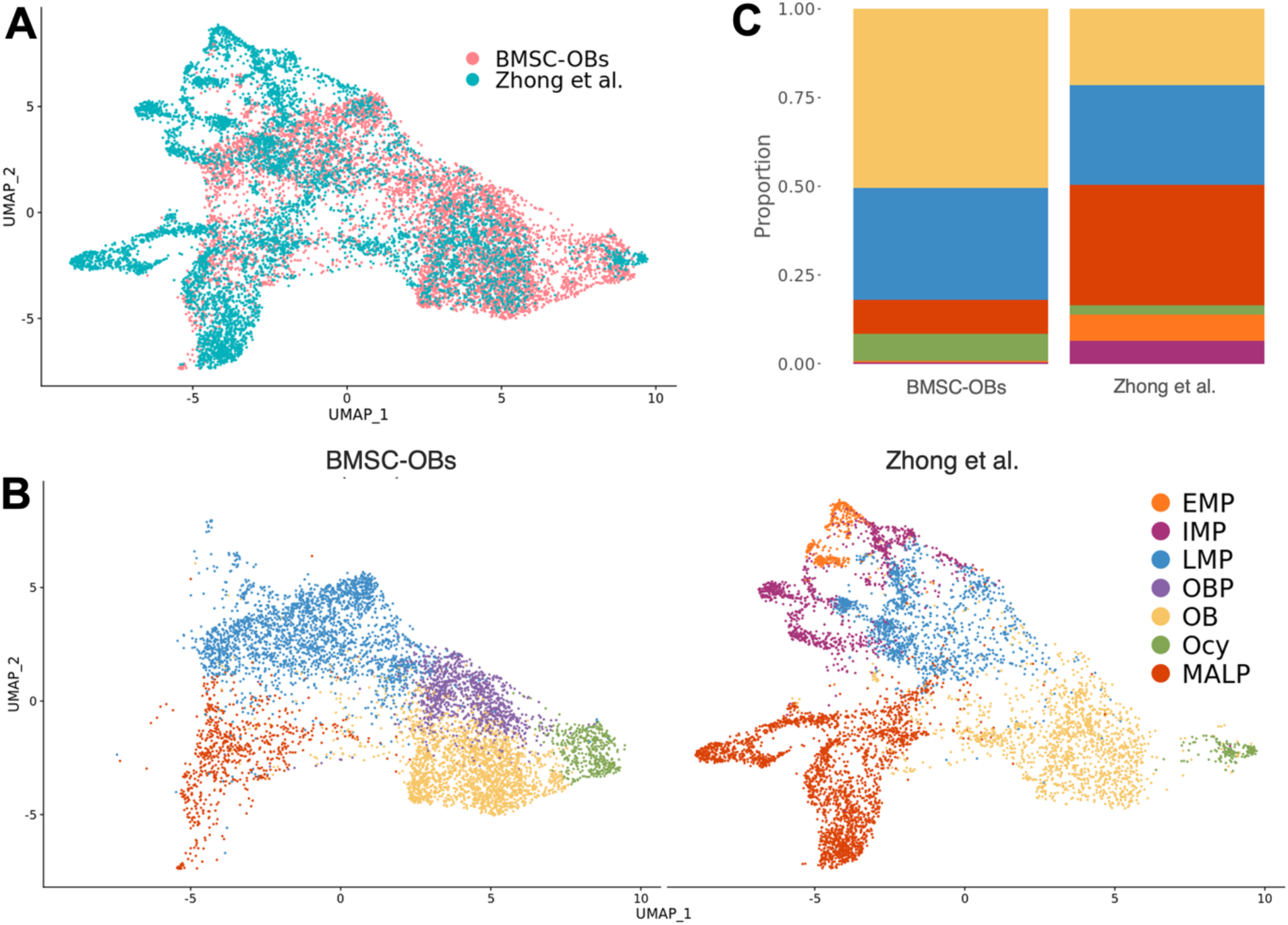
ScRNA-seq of BMSC-OB and scRNA-seq data derived from cells harvested *in vivo* cluster similarity and are transcriptomically identical. **A)** Single cells in UMAP space after integration of both the BMSC-OBs and Zhong et al. (2020) scRNA-seq datasets via Canonical Correlation Analysis (CCA). **B)** UMAPs with cell cluster annotation of the integrated data split based on the dataset. **C)** Stacked bar chart representing the proportion of each cell type for each dataset.

### Transcriptomic profiles from scRNA-seq for individual cell-types are robust

One of our goals for future experiments will be to generate expression profiles for multiple mesenchymal cell-types in large populations of mice (or humans) for use in downstream applications such as eQTL analysis or the generation of networks to inform human GWAS. To evaluate how well cell-type specific expression profiles from scRNA-seq align with profiles generated via traditional bulk RNA-seq, we performed a correlation between the expression profiles derived from each of the six defined cell-types (five mesenchymal + one grouped immune cell cluster) examined in this study to bulk RNA-seq data (derived from psc-bulk data described above). We generated a “pseudo-bulk” profile (PB) from the scRNA-seq data by aggregating counts across all cells belonging to a specific cell-type to simulate a dataset representative of one derived from bulk sequencing methods. A high correlation was observed between both the bulk/psc-bulk profiles and the PB profile generated for the entire scRNA-seq dataset (r = 0.84 and r = 0.85, respectively; P<2.2×10^−16^) (**Figure 4A**). Upon comparison of individual PB profiles generated for each cluster in the scRNA-seq to the psc-bulk samples, PB profiles were highly correlated; for example, correlations of PB profiles for the osteoblast (OB) and osteocyte (Ocy) cell clusters were r = 0.83 (P<2.2×10^−16^) and r = 0.81 (P<2.2×10^−16^), respectively (**Figure 4B**). As expected, correlations were slightly higher when comparing PB vs. psc-bulk (rather than PB vs. bulk), likely due to the single-cell isolation procedure performed in the psc-bulk samples.

**Figure 4:**
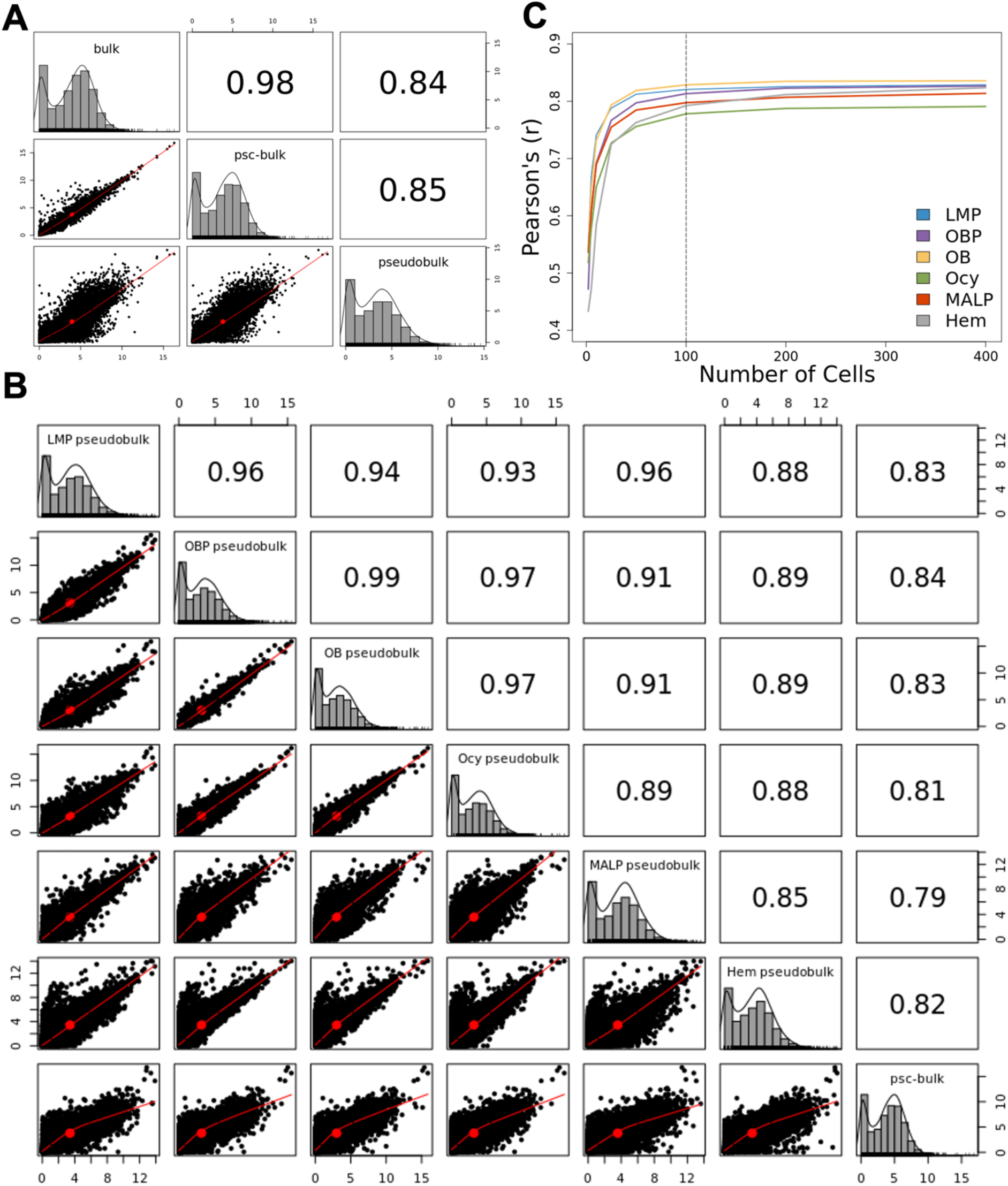
Transcriptomic profiles of individual cell types from scRNA-seq of BMSC-OBs are robust and representative of bulk RNA-seq data. **A)** Correlation between the counts per million (CPM) values from RNA counts derived for bulk, pooled single cell-bulk (psc-bulk), and pseudobulk (PB) samples. **B)** Correlation between the CPM values derived from the psc-bulk and PB profiles for each annotated cell cluster of the BMSC-OB scRNA-seq data. **C)** Correlation between overall PB and PBs for each cell-type generating using different numbers of cells (2 to 400).

Additionally, we estimated the minimum number of cells per cluster required to generate robust cell-type expression profiles by randomly selecting from 2 to 400 cells from each cluster, generating a PB profile (as described above), and subsequently calculating the correlation between each cell-type PB profile to the psc-bulk sample. Calculated correlations plateaued for all cell-types at ∼100 cells (**Figure 4C**). These data indicate that aggregated data across at least 100 cells from a given cell-type approximates data generated from bulk RNA-seq.

### Frequency of osteogenic cell-types are highly variable across DO mice

Because the BMSC-OB scRNA-seq dataset consists of multiple samples pooled into one, we used Souporcell^31^ genotype deconvolution to assign a mouse of origin to each cell with high confidence (**Supplemental Data 4**). Five genotypically distinct clusters (genotypes) were inferred by Souporcell from the scRNA-seq data based on SNPs captured in the sequenced cDNA. Genotype clusters were assigned to their corresponding DO mouse ID by comparing allele calls made by the variants captured between Souporcell and genotypes previously generated on all five DO mice using the GigaMUGA genotyping^16,32^. Of the 67,056 total variants identified by Souporcell, 0.87% (581) were also captured by the GigaMUGA arrays (143,259 total). DO mouse IDs were assigned based on the highest percentage of matching allele calls made upon pairwise comparison between Souporcell cluster and GigaMUGA arrays. After assigning a mouse of origin for all cells in the scRNA-seq data, we quantified differences in the frequencies of various cell-types contributed by each mouse (**Figure 5A**). For example, mouse #50 had a higher frequency of LMPs and MALPs and fewer osteoblasts and osteocytes compared to the other four mice (**Figure 5A, 5B**). Pooling samples for scRNA-seq, coupled with genotype deconvolution downstream, is an approach that is scalable for multi-sample input, which is necessary to perform population-level investigations.

**Figure 5:**
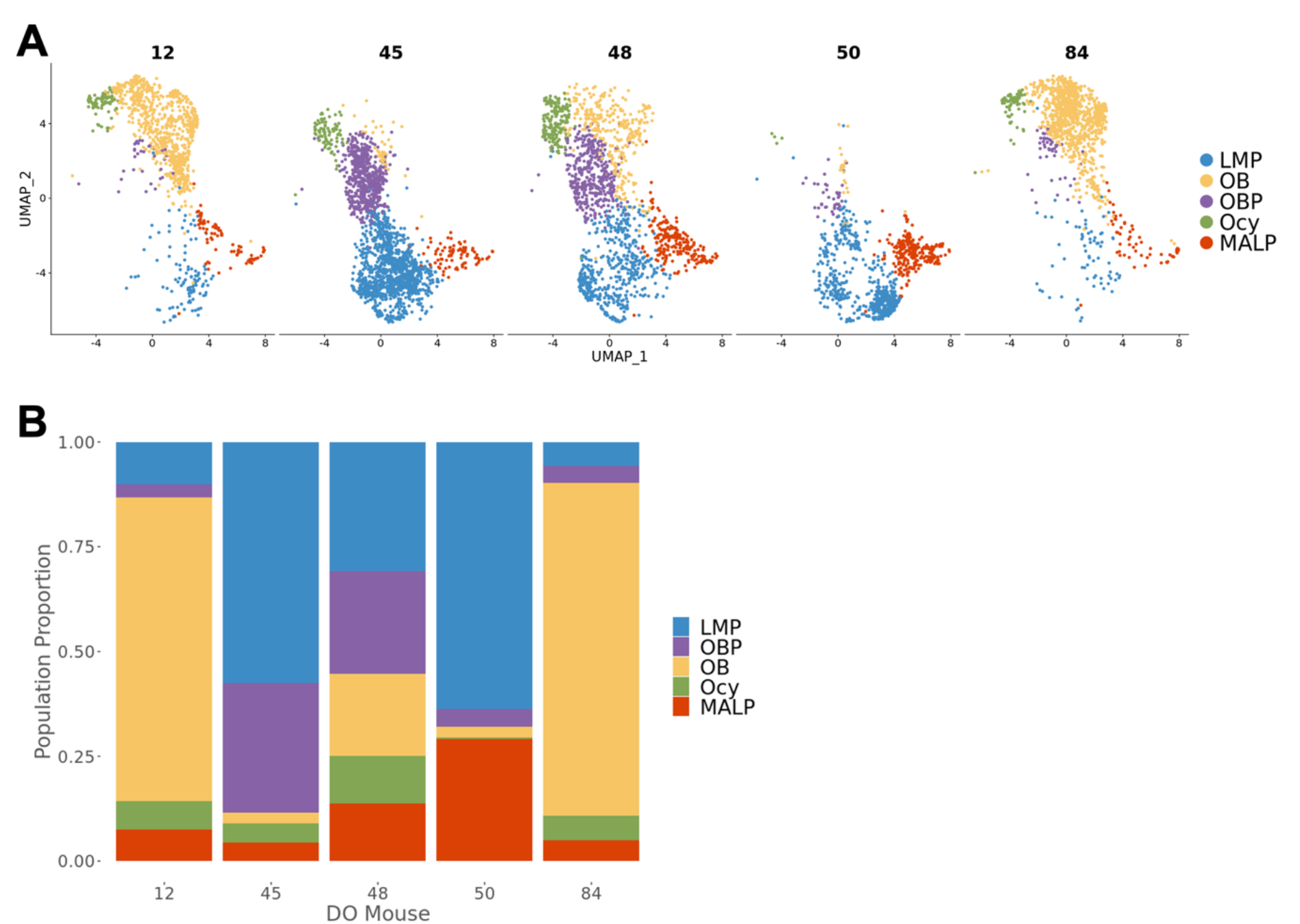
Cell type frequencies captured by scRNA-seq are highly variable across individual DO mice. **A)** UMAPs of the BMSC-OBs derived from the five DO mice, split based on each mouse (12, 45, 48, 50, 84). **B)** Stacked bar chart representing the proportion of each cell type derived from each mouse.

### BMSC-OBs show expected gene regulatory networks

Cell-type identification is largely based on the association of canonical and highly expressed genes with certain cell-types; however, underlying gene regulatory networks (GRNs) provide insight into how expression is coordinated. Moreover, GRN inference can be used to establish gene expression profiles for cell-types of interest by elucidating which specific combinations of transcription factors (TFs) are responsible for the expression of downstream target genes. We used SCENIC^33^ to better understand the GRNs that characterize the cell states in BMSC-OBs. The SCENIC analysis pipeline first generates regulatory modules inferred from co-expression patterns, which are used to form “regulons” consisting of a core TF that governs the expression of predicted target genes. Next, target genes are pruned based on enrichment of the TF cis-regulatory motifs located upstream or downstream of target genes in the regulon (**Supplemental Data 5)**. Lastly, the activity of regulons is quantified across individual cells (**Supplemental Data 6)**.

We applied the SCENIC analysis pipeline to the BMSC-OBs and resolved distinct regulons associated with each cell cluster in the BMSC-OB dataset (**Figure 6**). Regulons were robust in activity (**Figure 6A, 6B, Supplemental Data 7**) and specific for each cell-type (**Figure 6C, 6D, Supplemental Data 8**). For example, *Sp7* (Osterix), a key TF known to be involved in osteoblast differentiation, was found to be more specifically associated and highly active in the OBP cell cluster (**Figure 6C, 6D)**. Similarly, we show *Pparg* is a highly active regulon and exclusively associated with MALPs (**Figure 6C, 6D)**, consistent with its role as a master regulator of adipogenesis. This analysis suggested that not only do BMSC-OB cell-types show similar transcriptomic signatures to the same cells isolated directly from bone, but cell circuits (i.e., GRNs) are also similar.

**Figure 6:**
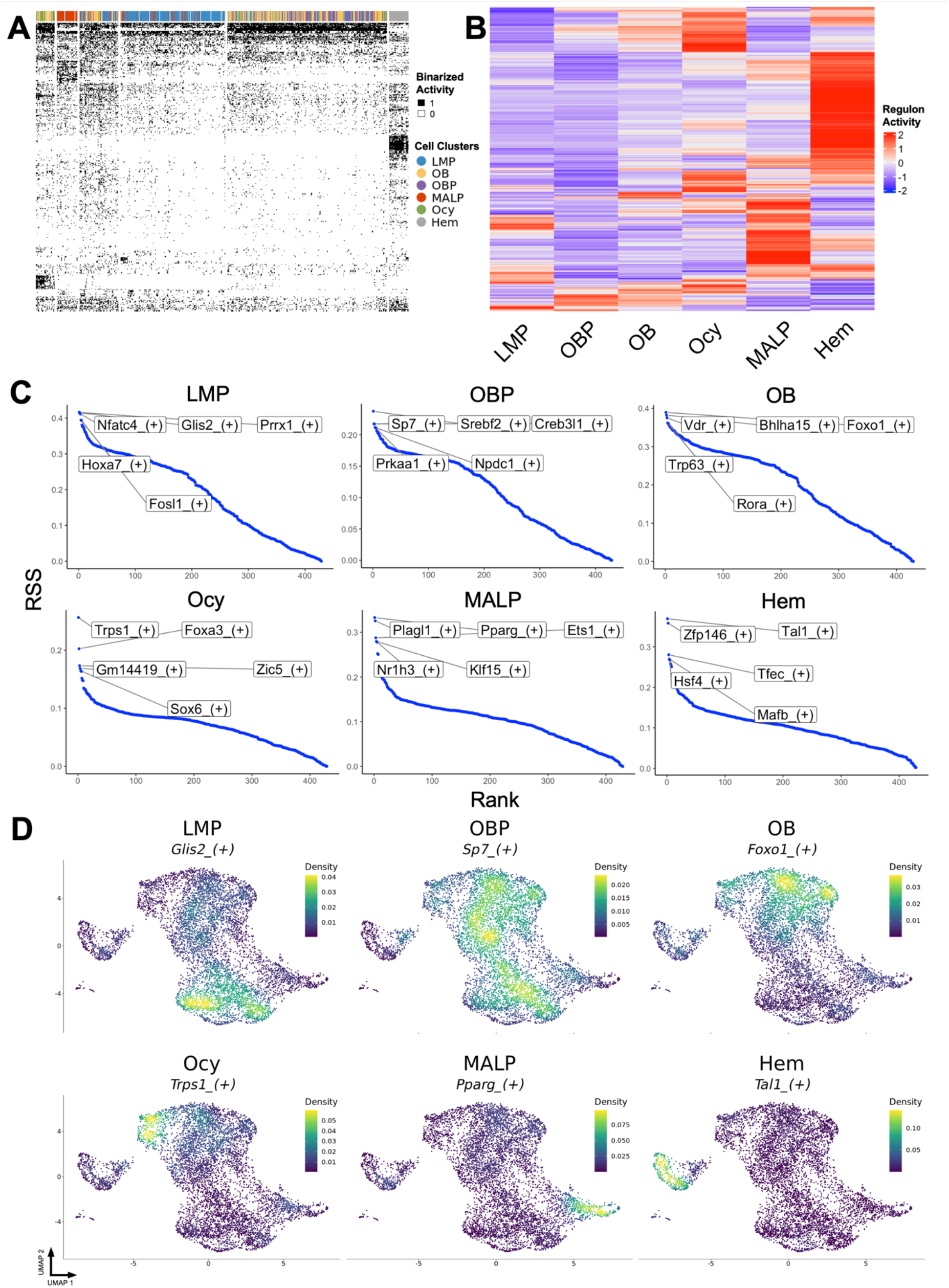
SCENIC gene regulatory network (GRN) analysis reveals expected transcriptomic activity and validates the identities of cell-types in BMSC-OBs. **A)** Binarized heatmap SCENIC regulon activity results, where “1” indicates active regulons; “0” indicates inactive regulons. **B)** Heatmap of SCENIC results portraying the scaled average for regulon activity in each annotated cell cluster, where the color key from blue to red indicates activity levels from low to high, respectively. **C)** Plots of the top five regulons with the highest specificity score (RSS) for each cell cluster. **D)** Density plots portraying the regulon-weighted 2D kernel density of select regulons for each cell cluster.

### MALPs and osteogenic cells capture BMD heritability identified by GWAS

We next used CELLECT^34^ to evaluate the relevance of identified cell-types with regards to mediating the effects of GWAS. CELLECT integrates disease heritability estimates with cell-type expression specificity from scRNA-seq data to identify cell-types that capture a significant component of the heritability for a disease or trait. We applied CELLECT to the cell-types identified in BMSC-OBs and those identified by Zhong et al. (2020). We observed that genes with selective expression in MALPs, OBs, and Ocys from both datasets were significantly (P < 0.05) enriched for BMD heritability. In addition, IMPs and LMPs in the Zhong et al. (2020) dataset were also significant. Non-mesenchymal lineage cells, which are mostly immune cells in both datasets were not significant **(Table 1**). Interestingly, osteoclasts captured in Zhong et al. (2020) dataset were not identified as significant in the CELLECT analysis (**Table 1**).

**Table 1:** CELLECT cell-type prioritization on all cell-types annotated in the BMSC-OBs and Zhong et al. (2020) scRNA-seq datasets.

| scRNA-seq dataset | Cell type | Beta | Beta SE | P-value |
| --- | --- | --- | --- | --- |
| Zhong et al. | MALP | $5.88 \times 10^{-8}$ | $1.84 \times 10^{-8}$ | $6.92 \times 10^{-4}$ |
| Zhong et al. | OB | $4.80 \times 10^{-8}$ | $1.56 \times 10^{-8}$ | $1.05 \times 10^{-3}$ |
| Zhong et al. | Ocy | $5.91 \times 10^{-8}$ | $2.15 \times 10^{-8}$ | $3.03 \times 10^{-3}$ |
| BMSC-OBs | Ocy | $5.70 \times 10^{-8}$ | $2.16 \times 10^{-8}$ | $4.18 \times 10^{-3}$ |
| BMSC-OBs | MALP | $4.86 \times 10^{-8}$ | $1.86 \times 10^{-8}$ | $4.57 \times 10^{-3}$ |
| Zhong et al. | IMP | $3.61 \times 10^{-8}$ | $1.68 \times 10^{-8}$ | $1.57 \times 10^{-2}$ |
| Zhong et al. | LMP | $3.09 \times 10^{-8}$ | $1.71 \times 10^{-8}$ | $3.55 \times 10^{-2}$ |
| BMSC-OBs | OB | $6.24 \times 10^{-8}$ | $3.56 \times 10^{-8}$ | $3.98 \times 10^{-2}$ |
| Zhong et al. | EMP | $2.86 \times 10^{-8}$ | $1.79 \times 10^{-8}$ | $5.52 \times 10^{-2}$ |
| Zhong et al. | CH | $1.96 \times 10^{-8}$ | $1.38 \times 10^{-8}$ | $7.81 \times 10^{-2}$ |
| Zhong et al. | Mural | $9.12 \times 10^{-9}$ | $1.66 \times 10^{-8}$ | $2.91 \times 10^{-2}$ |
| BMSC-OBs | LMP | $-4.57 \times 10^{-9}$ | $2.04 \times 10^{-8}$ | $5.89 \times 10^{-2}$ |
| BMSC-OBs | OBP | $-5.85 \times 10^{-9}$ | $1.77 \times 10^{-8}$ | $6.30 \times 10^{-1}$ |
| Zhong et al. | Erythrocyte | $-8.07 \times 10^{-9}$ | $1.73 \times 10^{-8}$ | $6.79 \times 10^{-1}$ |
| Zhong et al. | Mono | $-3.03 \times 10^{-8}$ | $1.60 \times 10^{-8}$ | $9.71 \times 10^{-1}$ |
| Zhong et al. | MF | $-2.98 \times 10^{-8}$ | $1.52 \times 10^{-8}$ | $9.75 \times 10^{-1}$ |
| Zhong et al. | EC | $-2.20 \times 10^{-8}$ | $1.10 \times 10^{-8}$ | $9.77 \times 10^{-1}$ |
| Zhong et al. | B-cell | $-3.47 \times 10^{-8}$ | $1.63 \times 10^{-8}$ | $9.83 \times 10^{-1}$ |
| Zhong et al. | OC | $-4.66 \times 10^{-8}$ | $1.55 \times 10^{-8}$ | 1 |
| Zhong et al. | Granulo | $-3.56 \times 10^{-8}$ | $9.95 \times 10^{-9}$ | 1 |
| BMSC-OBs | Hem | $-5.90 \times 10^{-8}$ | $1.36 \times 10^{-8}$ | 1 |
| Zhong et al. | T-cell | $-6.45 \times 10^{-8}$ | $1.35 \times 10^{-8}$ | 1 |
| Zhong et al. | HSC | $-6.28 \times 10^{-8}$ | $1.28 \times 10^{-8}$ | 1 |
| Zhong et al. | GP | $-5.44 \times 10^{-8}$ | $1.11 \times 10^{-8}$ | 1 |

Beta is regression effect size estimate for given annotation. Beta SE is the standard error for the regression coefficient. P-value is the one-sided test ( $\beta > 0$ ) association between BMD GWAS signal heritability and each annotated cell-type. P-values $< 0.05$ are highlighted in red. Cell-type abbreviations: OB: osteoblast; OBP: osteoblast progenitor; Ocy: osteocyte; EMP: early mesenchymal progenitor; IMP: intermediate mesenchymal progenitor; LMP: late mesenchymal progenitor; MALP: marrow adipogenic lineage precursors; CH: chondrocyte; HSC:
hematopoietic stem cell; EC: endothelial cell; GP: granulocyte progenitor; Hem: Hematopoietic lineage cells; OC: osteoclast; Granulo: granulocyte; MF: macrophage; Mono: monocyte; Mural: mural cells; T-cell: T-lymphocyte; B-cell: B-lymphocyte.

## Discussion

A considerable challenge faced upon analyzing GWAS is identifying the causal genes impacted by significant associations. Integrating transcriptomics data has proven invaluable for accomplishing this goal. Colocalizing genetic variation impacting gene expression with GWAS associations can identify putative causal genes influencing disease. Moreover, integrating single-cell transcriptomics data can provide the cellular context in which causal genes are most likely to be impactful. In the context of osteoporosis research, the generation of population-scale transcriptomics data at single-cell resolution would aid in gene discovery. Here, we demonstrate the use of BMSCs cultured under osteogenic conditions (BMSC-OBs) from the Diversity Outbred (DO) mouse population coupled with scRNA-seq can serve as a model to generate single-cell transcriptomics data of mesenchymal cell-types relevant to bone. We show that after subsequent culturing under osteoblast differentiation, there was an enrichment in the relative frequencies of osteoblasts and osteocyte-like cells, compared to cells isolated *in vivo* using a mesenchymal lineage reporter. Additionally, the model yields adipogenic progenitor cells and their transcriptomic signature is nearly identical to the MALPs identified in Zhong et al. (2020). These cells are classified as a stable intermediary cell-type along the adipogenic differentiation route after mesenchymal progenitors and before more mature, lipid-laden adipocytes (LiLAs)^23^. Thus, BMSC-OBs contain many of the key mesenchymal cell-types and leads to an enrichment of osteoblasts and osteocyte-like cells that we demonstrated were to likely be the most relevant for informing GWAS.

We addressed the technical challenges posed with our approach, such as the single-cell isolation procedure used to liberate BMSC-OBs from a highly mineralized matrix *in vitro*. This procedure consists of an approximately two-hour process involving incubations with proteases and EDTA, raising the concern of technical effects impacting the integrity and quality of the isolated cells for scRNA-seq. In the bulk vs. psc-bulk experiment, we sought to characterize the impact of the single-cell isolation procedure on gene expression. Despite the induction of inflammation/stress-related genes in the psc-bulk sample, the overall gene expression profiles between bulk and psc-bulk samples were highly correlated and any observed change in gene expression had a negligible impact on global transcriptomic signatures or downstream annotation of mesenchymal cell-types. However, care should be taken when interpreting the expression of individual genes, especially those identified to be responsive to the isolation procedure.

We also assessed the biological informativeness of BMSC-OBs by comparing them to the same cells isolated directly from bone. Upon comparison of both scRNA-seq datasets, we found that the transcriptomic signatures of BMSC-OB cell-types did not differ compared to the cells isolated by Zhong et al. (2020). Nevertheless, differences between the two datasets were observed, namely the absence of early/intermediate mesenchymal progenitor (EMP, IMP) populations in the BMSC-OB dataset, which is likely due to the maturation of LMPs beyond EMP/IMP cell stages during the *in vitro* osteoblast differentiation. Importantly, these results indicate that individual cell-types in BMSC-OBs are nearly identical, in the context of transcriptomic signatures, to their counterparts in bone.

A number of approaches have been used to profile individual bone cells. These include scRNA-seq on whole bone marrow^22,35^, using fluorescence-activated cell sorting (FACS) on marrow to enrich for mesenchymal lineage cells^36^, the digestion of bone combined with FACS^37^, and FACS in lineage specific reporter mice^38^. These studies have provided important insights into the cellular landscape of bone and the identity of skeletal stem cells. However, none of these approaches were developed with the goal of investigating bone cells at the population-scale in mice or humans. These approaches isolate a wide range of cells, many of which provide little insight in the context of informing BMD GWAS. Profiling non-relevant cells significantly increases cost and makes population screening less feasible. As an alternative, BMSC-OBs have several attractive attributes. First, it is simple, marrow is relatively easy to isolate from populations of mice, or even humans, and isolating BMSCs based on plastic adherence is cost-effective and straightforward. Second, we show that osteoblasts and osteocyte-like cells are some of the most relevant to BMD GWAS and we are able to enrich for these cells by culturing under osteogenic conditions. Third, as we have shown, there are few transcriptomic differences in cultured cells as compared to cells isolated directly from mice *in vivo*. Fourth, we do not need to use FACS or specific reporter mice, making it possible to perform this approach in any population of mice, and potentially humans. Fifth, because we enriched for key cell-types, the number of cells to sequence is lower.

A valid concern of population-based scRNA-seq studies is cost associated with increasing scalability and sample throughput. Using BMSC-OBs, we remedy this challenge by pooling single cells derived from multiple mice into a single sample for scRNA-seq. Because each DO mouse is genotypically distinct from one another, we performed a genotype deconvolution on the scRNA-seq data downstream. We were able to associate a “mouse-of-origin” to each single cell derived from our cohort of DO mice. Genotype deconvolution enables other downstream analyses, such as quantifying differences in certain cell fractions between samples. While the sample size of our mouse cohort in this study was small (N = 5), we observed significant differences in cell-type frequencies between our mice. These differences likely reflect variation in cell-type composition of the starting BMSCs and differences in the rate/efficiency of osteoblast differentiation arising as a function of mouse-specific genotype and environmental effects. With this study serving as a proof-of-concept, BMSC-OBs feasibly permits scalability and increased sample throughput, which is necessary to inform GWAS.

One of the major limitations of our approach is that the BMSC-OB model does not capture all cell-types relevant to bone. For example, it does not capture osteoclasts. However, it is important to note that in our CELLECT analysis BMD heritability was not enriched in genes whose expression was more specific to osteoclasts from the Zhong et al. (2020) dataset. It is unclear why osteoclasts were not significant and may be due to cross-sectional measures of BMD being more so a product of peak bone mass and osteoblast-mediated bone accrual, than bone loss, a process driven by osteoclasts, or the fact that these were likely immature osteoclasts as mature cells would be too large to be captured for sequencing. Although, marrow adiposity^39^ and MALPs^23^ have been demonstrated to significantly influence bone mass, it was somewhat surprising that the CELLECT analysis identified a significant association between gene expression specificity in MALPs and BMD heritability. This potentially suggests that many BMD GWAS associations impact genes regulating BMD via MALPs. Future studies should seek to identify specific associations that may be working through marrow adipocytes.

Here, we described how the osteogenic differentiation of BMSCs can facilitate the generation of large-scale scRNA-seq data for mesenchymal lineage cells derived from the DO mouse population. Based on findings gathered here, the transcriptomic profiles generated from BMSC-OBs will serve as a valuable biological input for future genetic analysis. For example, cell-type specific, co-expression networks can be used as input to perform directed Bayesian network reconstruction and Key Driver Analysis (KDA), as previously described in^16^. These subsequent analyses can aid in informing GWAS and highlighting putatively novel genes driving disease. We have demonstrated that the BMSC-OB model has the potential to facilitate more holistic genotype-to-phenotype investigations, which will aid in our understanding of the genetics of bone mass and lead to the identification of novel therapeutic targets that could be targeted to treat and prevent osteoporosis.

## Methods

### Sample preparation and *in vitro* cell culture of BMSCs

Bone marrow extraction and subsequent cell culture was performed as described in^16^. In brief, the left femur was isolated and cleaned thoroughly of all muscle tissue followed by removal of its distal epiphysis. Marrow was exuded by centrifugation at 2000×g for 30 s into a sterile tube containing 35 μL fetal bovine serum (FBS, Atlantic Biologicals). The marrow was then triturated 6 times on ice after addition of 150 μL of cold freezing media (90% FBS, 10% Dimethyl Sulfoxide (DMSO, Fisher). Marrow was placed into a Mr. Frosty Freezing Container (Nalgene) for the purpose of slow cooling, stored overnight at -80°C, and transferred to liquid nitrogen for long-term storage. In preparation for cell culture, samples were thawed at 37°C, resuspended in 5 mL bone marrow growth media (MEM alpha, Gibco), 10% FBS, 1% Penicillin Streptomycin (Pen/Strep, Gibco), 1% Glutamax (Gibco), and then subjected to red blood cell lysis by resuspending with 5 mL 0.2% NaCl for 20 s then thorough mixing of 1.6% NaCl. Cells were pelleted, resuspended into 1 mL bone marrow growth media, and cultured in 48 well tissue culture plates. Samples were incubated in a 37°C, 5% CO_2_, 100% humidity incubator and left undisturbed for 3 days. Thereafter, media was aspirated and replaced daily. After 6 days, cells were washed and then underwent a standard *in vitro* osteoblast differentiation protocol for 10 days by replacing bone marrow growth media with 300 μL osteogenic differentiation media (Alpha MEM, 10% FBS, 1% Pen/Strep, 1% Glutamax, 50 μg/mL Ascorbic Acid (Sigma), 10 mM B-glycerophosphate (Sigma), 10 nM Dexamethasome (Sigma).

### Single-cell isolation procedure

The isolation procedure outlined below was inspired by^40^. Mineralized cultures were washed twice with Dulbecco’s Phosphate Buffered Saline (DPBS, Gibco). 0.5 mL of 60 mM Ethylenediaminetetraacetic acid, pH 7.4 made in DPBS (EDTA, Fisher) was added and cultures were incubated at room temperature (RT) for 15 min. EDTA solution was aspirated, replaced, and cultures were incubated again at RT for 15 min. Cultures were then washed with 0.5 mL Hank’s Balanced Salt Solution (HBSS, Gibco) and incubated with 0.5 mL 8 mg/mL collagenase (Gibco) in HBSS/4 mM CaCl_2_ (Fisher) for 10 min at 37°C with shaking. Cultures were triturated 10 times and incubated for an additional 20 min at 37°C. Cultures were then transferred to a 1.5 mL Eppendorf tube and centrifuged at 500×g for 5 min at RT. Cultures were resuspended in 0.5 mL 0.25% trypsin-EDTA (Gibco) and incubated for 15 min at 37°C. Cultures were then triturated and incubated for an additional 15 min, after which 0.5 mL of media was added, triturated, and spun at 500×g for 5 min at RT. Cultures were then resuspended in 0.5 mL osteogenic differentiation media and cells were counted.

### Bulk RNA-seq analysis

Total RNA was extracted using a RNeasy Micro Kit (QIAGEN) and poly-A selected RNA was sequenced via GENEWIZ (South Plainfield, NJ, USA). RNA-seq analysis was performed using a custom bioinformatics pipeline. Briefly, FastqQC^41^ and RSeQC^42^ were used to assess the quality of raw reads. Adapter trimming was completed using Trimmomatic^43^. Sequences were aligned to the GRCm38 reference genome^44^ using the SNP and splice aware aligner HISAT2^45^. Genome assembly and abundances in counts per million (CPM) were quantified using StringTie^46^. Differential expression analysis was performed using the DESeq2^47^ package in R.

### Single-cell analysis pipeline

After the single-cell isolation procedure, cells from all five mice were pooled and concentrated to 800 cells/μL in sterile PBS supplemented with 0.1% BSA. The single-cell suspension was loaded into a 10X Chromium Controller (10X Genomics, Pleasanton, CA, USA), aiming to capture 8000 cells, with the Single Cell 3’ v2 reagent kit, according to the manufacturer’s protocol. Following GEM capturing and lysis, cDNA was amplified (13 cycles) and the manufacturer’s protocol was followed to generate the sequencing library. The library was sequenced on the Illumina NextSeq500 and the raw sequencing data was processed using 10X Genomics Cell Ranger toolkit (version 5.0.0). The reads were mapped to the GRCm38 reference genome^44^. Overall, 8990 cells were sequenced, to a mean depth of 57,717 reads per cell. Sequencing data is available on GEO at accession code GSE152806.

Analysis of the scRNA-seq data was performed using Seurat^27^ (version 4.1.1). Features detected in at least three cells where at least 200 features were detected were used. We used Souporcell^31^ (described below) to remove doublet cells. We then filtered out cells with less than 800 reads and more than 5800 reads, as well as cells with 10% or more mitochondrial reads. This resulted in 7357 remaining cells. The resulting object underwent standard normalization, scaling, and the top 3000 features were modeled from the mean-variance relationship using Seurat’s “FindVariableFeatures” function. Cell-cycle markers identified by Tirosh et al. (2016) were regressed out using the “CellCycleScoring” and scaling functions. For subsequent dimensionality reduction, 14 principal components (PCs) were summarized, which was the number of PCs where the percent change in variation between the consecutive PCs was less than 0.1%. A kNN (k = 20) graph was created and cells were clustered using the Louvain algorithm at a resolution of 0.22. Cluster cell-types were manually annotated after performing differential gene expression analysis of each cell cluster relative to all other clusters using the Seurat “FindAllMarkers” function (**Supplementary Data 1**).

Trajectory inference/pseudotime analysis was performed using Slingshot^28^ (version 1.6.1) on osteogenic/adipogenic lineage cells with the starting cluster set as the LMPs. TradeSeq^48^ (version 1.4.0) was used to analyze gene expression along the trajectories by fitting a negative binomial generalized additive model (NB-GAM) to each gene using the “fitGAM” function with nknots = 8, which was determined by using the “evaluateK” function.

### Integration of datasets via Canonical Correlation Analysis (CCA)

CCA^30^ in Seurat was used to integrate *in vivo* scRNA-seq data derived from Zhong et al. (2020) (1.5 month and 3 month timepoints) with the BMSC-OB *in vitro* data. The Zhong et al. (2020) data was first pre-processed in the same fashion as the BMSC-OBs scRNA-seq dataset. Cell-types not present in the BMSC-OBs dataset were removed from the Zhong et al. (2020) data in order to portray only osteogenic and adipogenic lineage cells. After integration, the combined dataset was clustered and analyzed as described in the single-cell analysis pipeline (above).

### Souporcell

Upon performing Souporcell^31^ (version 2.0.0), barcoded cells identified as doublets were removed from the scRNA-seq count matrix during pre-processing of the data. Additionally, Souporcell was used to perform genotype deconvolution using the GRCm38 reference genome^44^. Five genotypically distinct clusters (genotypes) were inferred based on variants in the sequenced reads. Genotype clusters were assigned their corresponding DO mouse ID by comparing allele calls made by the shared variants captured between Souporcell and GIGA-MUGA arrays previously performed on all mice in the cohort. DO mouse IDs were assigned by making a pairwise comparison between each Souporcell genotype cluster and GigaMUGA array. The comparison yielding the highest percentage of matching allele calls indicated the identity/genotype of each mouse (**Supplemental Data 4**).

### SCENIC

pySCENIC (Single-Cell rEgulatory Network Inference and Clustering)^33^ (version 0.11.2) was used to infer gene regulatory networks. A fully-processed Seurat object containing cell-type annotations was transformed into a loom file by using SeuratDisk^49^ (version 0.0.0.9019). The loom file was subsequently used as input to the SCENIC workflow^50^. In brief, gene regulatory networks (GRNs) were built using GRNBoost^51^ to identify potential gene targets for each transcription factor (TF) based on co-expression. CisTarget^52^ was then used to select potential direct target genes of the governing TF of the co-expression modules. (**Supplemental Data 5**). The activity of the final regulons were calculated using AUCell^33^ (**Supplemental Data 6**). Regulon specificity score (RSS) is based on Jensen-Shannon divergence, as described in^53^ (**Supplemental Data 8**). Stable cell states were identified by analyzing the most active and specific regulons for each cluster as well as associated target genes.

### CELLECT

CELLECT^34^ (Cell-type Expression-specific integration for Complex Traits) (version 1.1.0) was used to identify likely etiologic cell-types underlying complex traits of both the BMSC-OBs and Zhong et al. (2020) datasets. CELLECT quantifies the association between the GWAS signal and cell-type expression specificity using the S-LDSC genetic prioritization model^54^. Summary statistics from the UK Biobank eBMD and Fracture GWAS^3^ (Data Release 2018) and cell-type annotations from each scRNA-seq dataset were used as input. Cell-type expression specificities were estimated using CELLEX^34^ (CELL-type EXpression-specificity) (version 1.2.1). The CELLECT output prioritizes likely etiologic cell-types for BMD (**Table 1**).

## Supporting information

Description of Supplemental Data

