## Supplementary material for "Evaluation of a scalable approach to generate cell-type specific transcriptomic profiles of mesenchymal lineage cells": Description of Supplemental Data

**List of Supplementary Tables**

**Table S1.** Differentially expressed genes (DEGs) for each cluster relative to all other clusters in the BMSC-OB scRNA-seq dataset

**Table S2.** DEGs analysis of bulk RNA-seq data from the psc-bulk vs. bulk experiment

**Table S3.** Gene Ontology (GO) enrichment analysis of the DEGs from the psc-bulk samples

**Table S4.** Souporcell genotype deconvolution results and percent accuracy of allele calls made in pairwise comparison between each Souporcell cluster and GIGAMUGA genotype microarray for each DO mouse.

**Table S5.** SCENIC results of the top five regulons based on Regulon Specificity Score (RSS) and associated targets genes for each regulon for each cell cluster

**Table S6.** SCENIC results portraying regulon activity for each regulon for each barcoded cell

**Table S7.** SCENIC results of scaled regulon activity for each cell cluster

**Table S8.** SCENIC results of the RSS scores for each regulon for each cell cluster
